# Ongoing HIV replication in lymph node sanctuary sites in treated patients contributes to the total latent HIV at a very slow rate

**DOI:** 10.1101/2023.02.18.529086

**Authors:** Sasan Paryad-zanjani, Aditya Jagarapu, Michael J. Piovoso, Ryan Zurakowski

## Abstract

Lymph nodes (LNs) serve as a sanctuary site for HIV viruses due to the heterogeneous distribution of the antiretrovirals (ARVs) inside the LNs. There is an ongoing debate whether this represents ongoing cycles of viral replication in the LNs or merely residual virus production by latently infected cells. Previous work has claimed that the measured levels of genetic variation in proviruses sampled from the blood were inconsistent with ongoing replication. However, it is not clear what rate of variation is consistent with ongoing replication in small sanctuary sites.

In this study, we used a spherically symmetric compartmental ODE model to track the HIV viral dynamics in the LN and predict the contribution of ongoing replication within the LN to the wholebody proviral pool in an ARV-suppressed patient. This model tracks the reaction-diffusion dynamics of uninfected, actively infected, and latently infected T-cells as well as free virus within the LN parenchyma and the blood, and distinguishes between latently infected cells created before ARV therapy and during ARV therapy.

We simulated suppressive therapy beginning in year 5 post-infection. Each LN sanctuary site had a volume of 1 ml, and we considered cases of 1ml, 30ml, and 250ml total volume, which represent a single active sanctuary site, moderate systemic involvement, and involvement of the total lymphoid tissue. Viral load in the blood rapidly dropped and remained below the limit of detection in all cases but remained high in the LN sanctuary sites. Novel latent cells increased systemically over time but very slowly, taking between 25 and 50 years to reach 5% of the total latent pool, depending on the volume of lymphoid tissue involvement.

Putative sanctuary sites in LNs are limited in volume and produce novel latent cells slowly. Assays to detect genetic drift due to such sites would require very deep sequencing if sampling only from the blood. Previous studies showing a lack of genetic drift are consistent with the expected contribution of ongoing replication in lymph node sanctuary sites.

## 1. Introduction

In 2020, 37.6 million people globally were living with HIV [1]. Combinatorial antiretroviral (ARV) therapy (cART) has been extremely successful in treating infected patients [12]. Nevertheless, there are still major challenges to eradicating HIV-1 cells. The cART cessation leads to an upsurge in viremia to a detectable level due to the activation of latently HIV-1 infected cells [9, 28]. Latent infected long-lived memory and naive CD4+ T-cells reside in reservoirs. These cells maintain the latent genome of HIV cells but do not produce viruses while in the resting phase. If activated by various stimuli, they can lead to viral rebound [10]. These reservoirs are seeded during the eclipse phase of the infection; hence it is challenging to start cART before the formation of reservoirs. However, early cART administration does lead to a reduction in the size of reservoirs [32]. Long-term follow-up studies on people living with HIV reveal that latent reservoirs decay slowly with a half-life of around 44 months, making eradication improbable [27, 8].

There is a controversial idea that lymph nodes serve as sanctuary sites in treated patients. Clinical studies have revealed that ARVs are distributed heterogeneously in some tissues. Many are excluded from patients’ lymph nodes (LNs) [20]. These LNs are potentially serving as sanctuary sites, as the low levels of ARV correlate with a high level of HIV RNA [13]. The existence of ongoing viral replication in sanctuary sites has been a long debate among scholars [21, 34, 11]. Raltegravir intensification studies used an alternative approach to detect ongoing replication, measuring changes in the 2-LTR circle frequency following raltegravir intensification in fully suppressed patients [4, 14]. Analysis of the dynamics of the measured 2-LTR concentration following intensification supported the conclusion that a fraction of the patients in each study had substantial ongoing replication, probably occurring in a drug sanctuary, prior to the raltegravir intensification [22]. Early studies such as [34] believed that the sequence changes of viral RNA in lymphocyte tissue are due to the ongoing residual replication. On the other hand, [17, 18] did not detect viral evolution by phylogenetic analysis in the samples sequenced from the HIV-1 patients’ blood. They demonstrated that despite the low levels of genetic sequence drift, the origin of the virus detected in the blood of treated patients did not result from newly infected cells but originated from cells, or the daughters of cells, that were already infected before treatment was initiated. In [3, 23], in order to investigate the possibility of ongoing replication in the lymph nodes, they sequenced the samples from both blood and lymph node tissues. They did not find any viral evolution, and their results were consistent with the clonal expansion of the cells before the initiation of ART. Our group has previously developed mathematical models of LNs as sanctuary sites [6]. In this study, we are looking at the effects of ongoing replication in sanctuary sites on the pool of latently infected cells. The current study shows that active replication in sanctuary sites is consistent with the low levels of observed genetic sequence drift by phylogenetic analysis seen in the previous studies.

Mathematical modeling has proven to help understand the HIV dynamics and drug effects in patients with AIDS. In an early study, Kepler and Perelson [19] proposed a mathematical model to capture the dynamic of virus production and the effects of the drug. Later, a model was proposed in [5], which was capable of simulating persistent and low-level replication in anatomical compartments due to the presence of latently infected cells in sanctuary sites. In [7], a model was developed in order to determine when we can cease the treatment without anticipating HIV relapse based on immune response strengths and latent reservoir size. In [6], a spatial, compartmental model of lymphoid follicles as sanctuary sites supporting ongoing viral replication was proposed. Different mathematical models to describe HIV persistence and latent reservoir formation are reviewed in [26].

In this study, a mathematical model is developed to describe the formation of the latent reservoirs in sanctuary sites such as lymph nodes in order to show why detecting genetic divergence using phylogenetic analysis is unlikely. We used a spatial model with ten compartments, where the first compartment represents the blood, and the following compartments represent different depths within a spherical sanctuary site with different diffusivity for ARVs. We show that ARV administration drops the virus and infected cells concentrations in the blood below the limit of detection. However, in the tissue, the concentrations remain above the limit of detection. Therefore, the model represents a sanctuary site that supports ongoing replication due to spatial isolation and low ARV concentration, as previously described [16]. We further show that it takes a significant amount of time for the latent reservoir cells formed after treatment, known as Novel Latent (NL) Cells, to become dominant over the latent reservoir cells formed before treatment, which are known as Archival Latent (AL) cells. We explore the relative abundance of the NL cells in circulation over time for several different sanctuary site sizes; even for very high levels of lymphoid tissue involvement, it takes many years for the NL cells to compose an appreciable fraction of the total latent cells in circulation. These results demonstrate why it is hard to capture NL cells after treatment starts in the bloodstream.

## 2. Methods

### Biological background

Lymph nodes have been proposed as a potential sanctuary site. Lymph nodes are extremely organized tissues. They are able to extract antigens from lymphoid to allow professional antigen-presenting cells (APCs) to present antigens to T-lymphocytes and initiate an appropriate immune response [30]. To make these functions possible, a highly isolated environment must be present that provides a proper amount of time for interactions between T-lymphocytes and APCs that lead to receptor-antigen recognition. The lobule is a parenchymal space highly isolated from blood and lymphoid, making these functions possible in the lymph nodes.

Lobules communicate with the rest of the body by the post-arterial vasculature and the lymphatic sinuses. The post-arterial vasculature in the lobule is composed primarily of High Endothelial Venules (HEVs) and non-fenestrated capillaries. HEVs have a very thick cellular boundary that allows only the active transport of most molecules. The reticular fibrous network forms a barrier around lobules that limits transport between the sub-capsular sinus network and excludes large particles [31]. The primary role of these barriers is to keep the isolation of the lobule’s antigen-enriched environment from the bloodstream and lymphatic [33]; however, this also results in a lack of drug penetration into the lobules. Direct imaging of antiretroviral drug distribution in macaque lymph nodes indicates variations of 10-30-fold in ARV concentration within the same lymph node [29]. IR-MALDESI imaging shows significant compartmentalization in the spatial distribution of drugs within lymph nodes. The concentration of drugs in sub-capsular and medullary sinuses is higher than in the lymph node lobules [29]. The lack of drugs in lobules results in an isolated environment, which provides an ideal site for virus and infected T-cell persistence.

### Description of the model

In this study, we used a set of ODEs to model the dynamics among uninfected target cells, *T*, infected target cells, *I*, Novel Latent (NL) infected target cells, *N*, Archival Latent (AL) infected target cells, *L*, and free virus *V*. We used an *M* compartmental model. The first compartment represents the blood, HEVs, and lymphatic vessels (*s = 1*). The remaining *M-1* compartments represent the different layers of a lymph node with different drug diffusivity (*s = 2:M*). These *M-1* compartments connect to the main compartment (*s* = 1), as figure 2 depicts.

**Figure 1:**
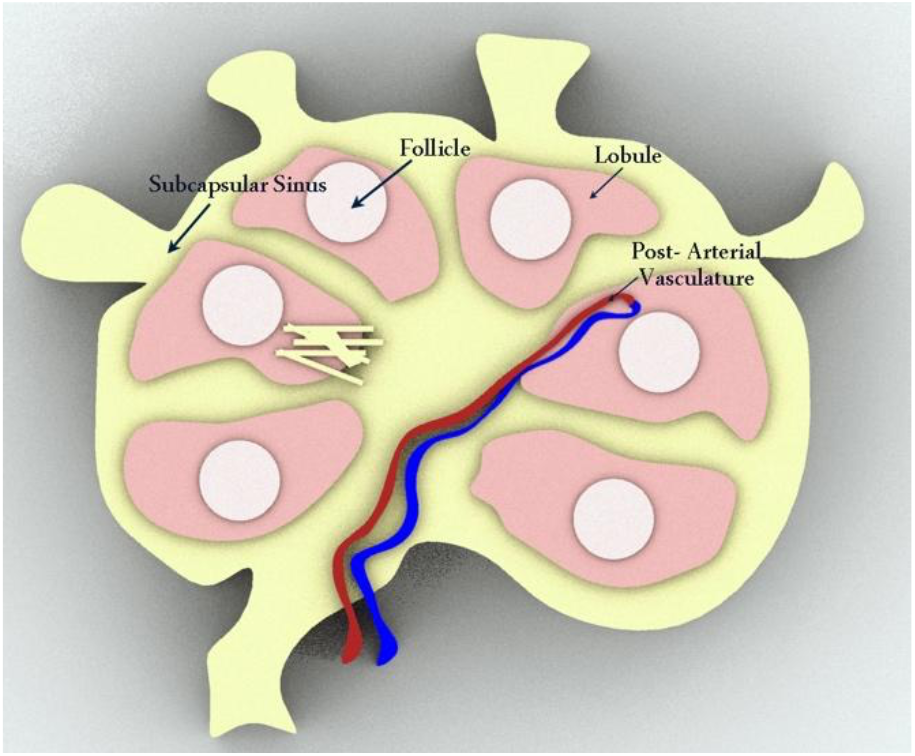
Lymph node

**Figure 2:**
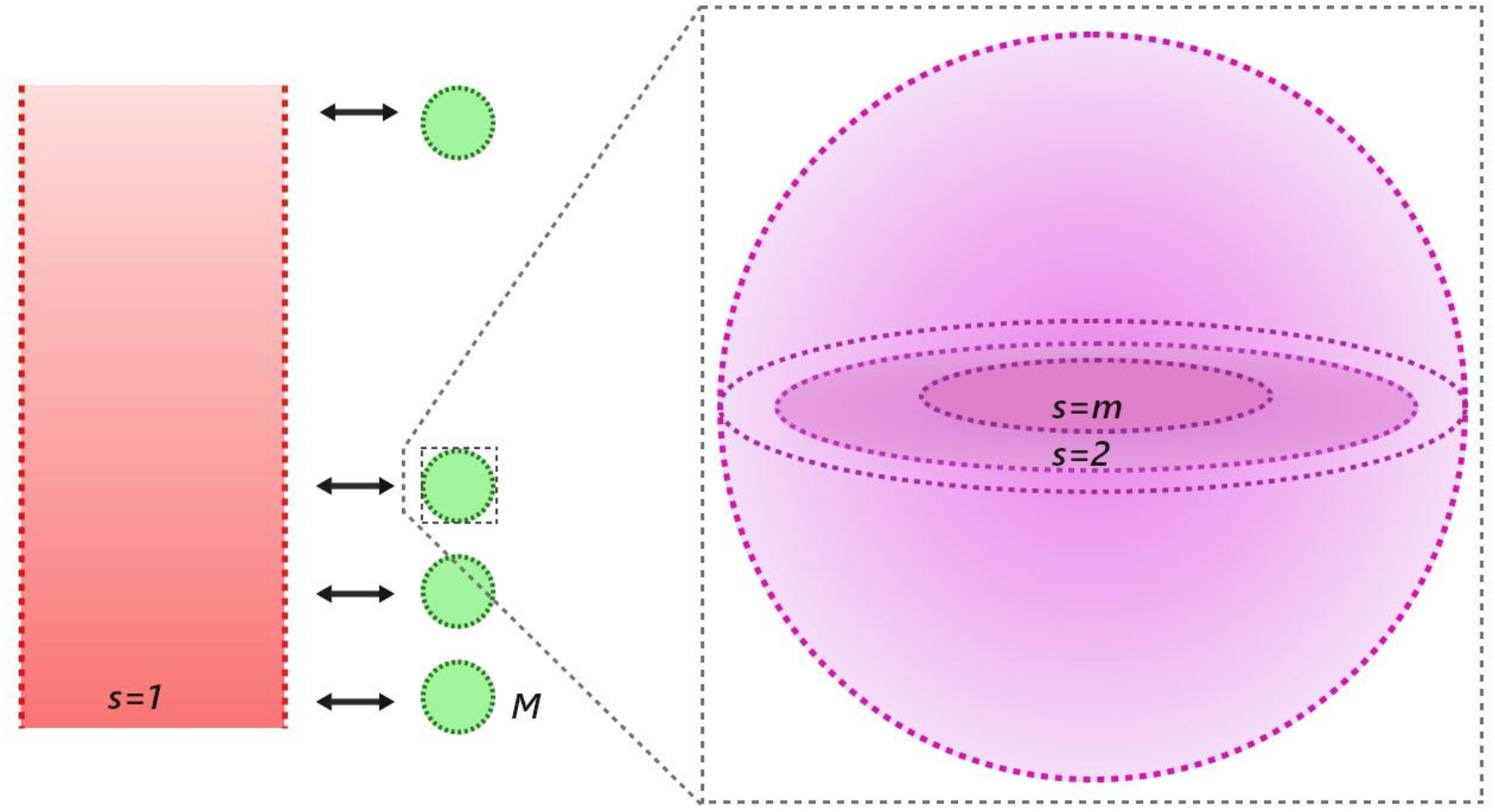
Configuration of compartments. On the left, the main compartment represents the blood, HEVs, and lymphatic vessels; in the middle, the m secondary spherical compartments represent the lymph nodes paracortex/follicle sites, and on the right the formulation of each sphere as M −1 concentric homogeneous shells.

In this model, we used the same assumptions as in [6]. First, T-cells can move inside the lymph node and from the inner sites of the lymph nodes to the blood and lymphatic vessels in a diffusion-like manner. Second, HIV is also transported inside the lymph node by diffusion. Third, HIV is only transported in or out of the lymphoid follicles inside infected T-cells [24]. Fourth, T-cells and virus motility are rapid enough such that the lymphoid follicle is well-described by concentric homogeneous spherical domains. Finally, we assume that a single, well-stirred compartment can jointly describe the blood and lymphatic vessels.

The first compartment representing the blood, HEVs, and lymphatic vessels has a connection with all of the lymph nodes through their outermost compartments, so it has a special form represented in equations 1–5. The equations 6–10 represent the remaining interior compartments. In equations 1 and 6, the T-cells, *T*, produced with a rate 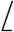, decay with a rate *d_T_T_s_*, and infected with rate *βT_s_v_s_*. The infection rate can be reduced by the activity of the drug with maximum efficacy of *η*. We hypothesize that the efficacy of the drug relies on the domain; hence, we include a spatially dependent drug distribution *θ_s_*. Infected cells, *I_s_*, decay with rate *d_I_,I_s_*, and the *α* fraction of infected cells become NL cells *M_s_*, or AL cells *L_s_* depending on the treatment phase. If the patient is in the OFF-treatment phase, *L_s_* are produced, and if the patient is in the ON treatment period, the *N_s_* is produced. Upon formation, the latent cells can proliferate with the rate *ρ*, decay with rate *d*, activate, and produce infected T-cells with rate *a*. The virus, *v_s_* is produced with rate *I_s_*, which can be reduced by activation of the drug. The virus also decays with rate, *d_v_*. Figure 3 shows the schematic of the reaction model that describes the interaction of viruses with drugs and immune systems. We described transportation among compartments as diffusion equations, where *M_s_* represents the set of the adjacent layers of *s*, *A_i,s_* represents the surface area between each layer in the sphere, represents the volume of the layer, and *D_Ti,s_*/*l, D_Ii,s_*/*l, D_Ni,s_*/*l, D_Li,s_*/*l*, and *D_vi,s_*/*l* represent the effective diffusivities of T-cells and HIV virus between layers. Equations 1–5 model the first compartment, which represents blood, HEVs, and lymphatic vessels.

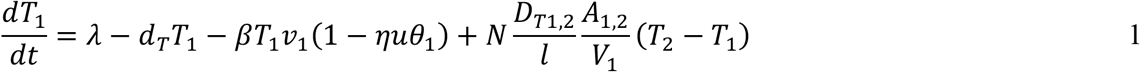

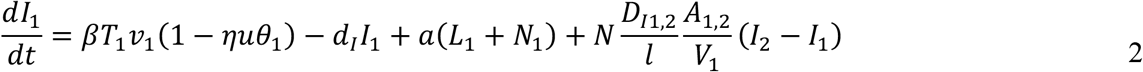

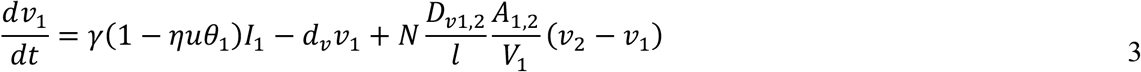

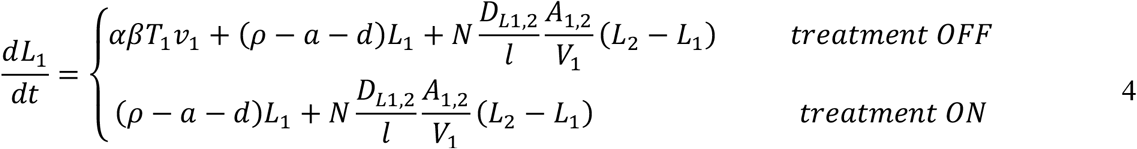

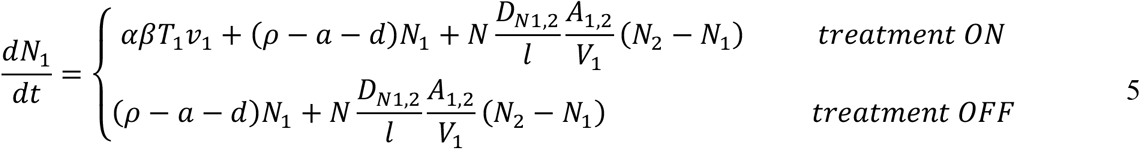

for the remaining *M-1* compartments, we have:

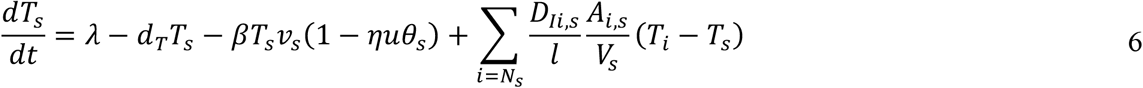

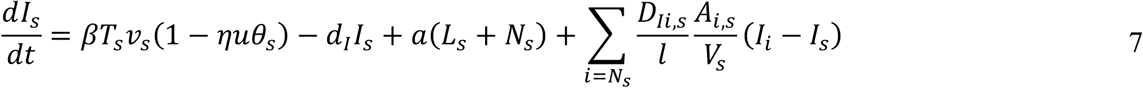

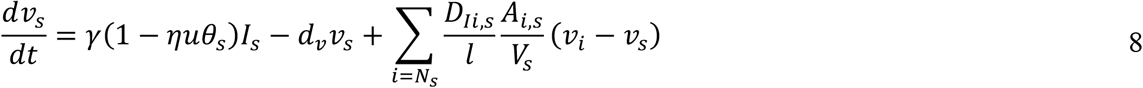

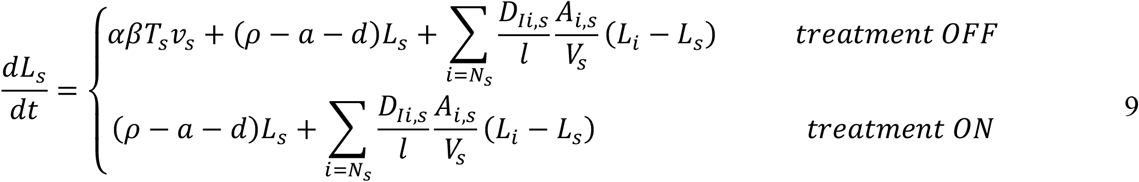

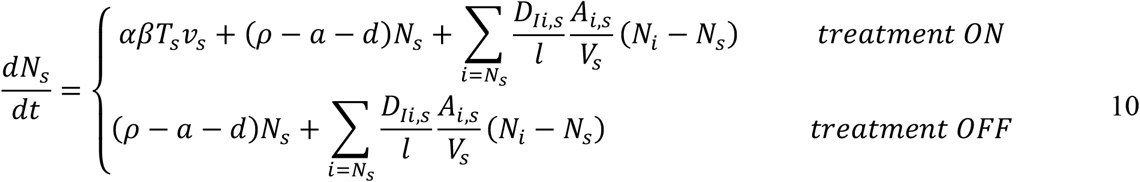

**Figure 3:**
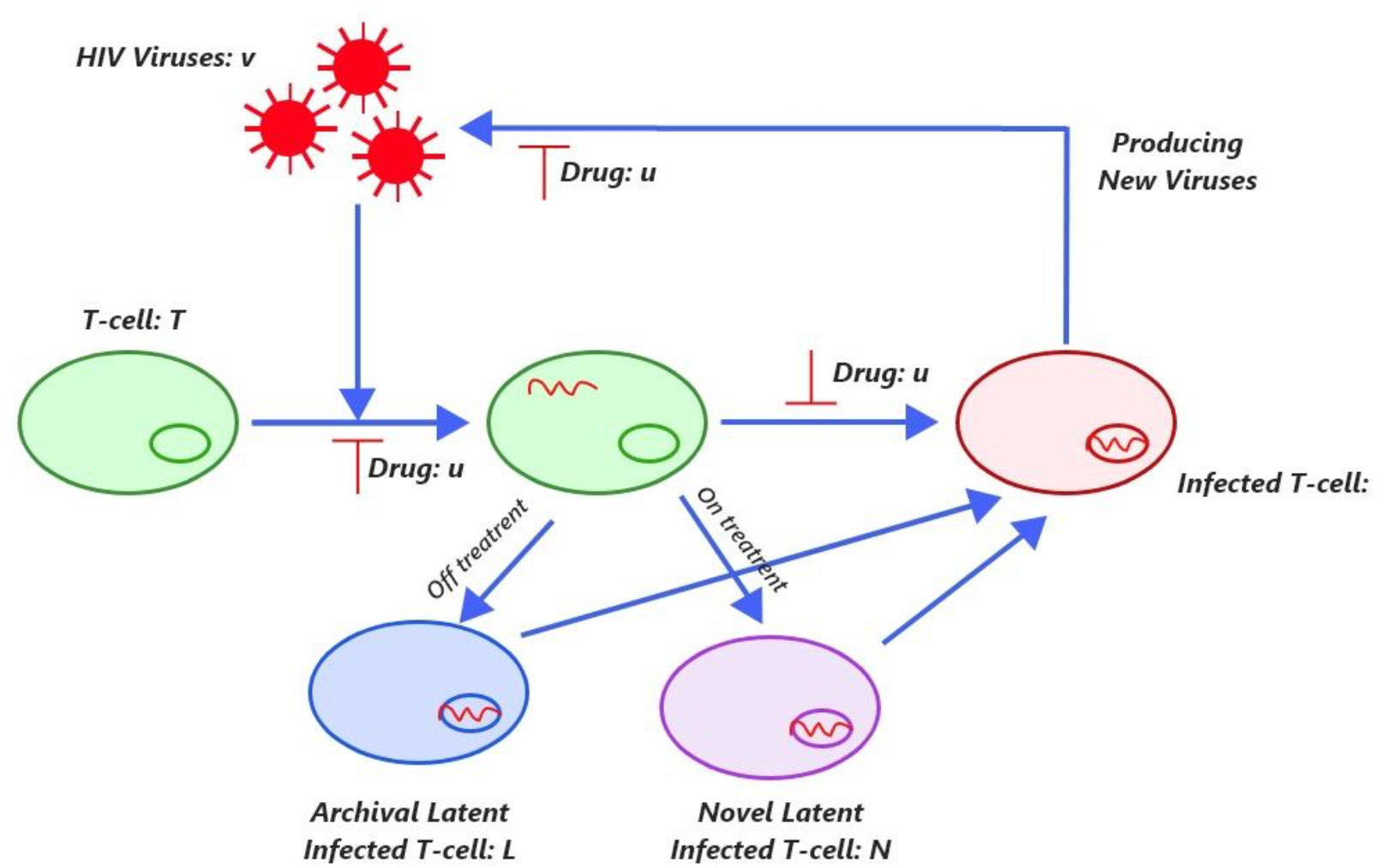
Illustration of the model. Description of the HIV interaction with immune system and drugs.

### Parameter values

We used data published in [22], and [15] to obtain the viral dynamic parameters and drug efficiency. [22] used a Bayesian Markov-Chain Monte Carlo technique to obtain the parameters from treatment interruption data taken from HIV patients who had 3–5 treatment interruption cycles each. These data produced a posterior distribution of parameter values. Table 1 shows the parameter ranges used in this study. The diffusion parameters depend on the size of the each compartment and the values for effective diffusivity of the T-cells and the virus. It was shown in [25] that hyperplastic lymphoid follicles can be as large as 1 mm in diameter. The relative permeability of T-cells and virus between the blood and follicle, are set to be 1/300, and 10^-8^ [mm/day] respectively [6]. T-Cells, and virus permeability inside the follicle are *0.1/1*, and 0.43/Z, respectively, where *I* = *r*/(*n* – 1) [6]. For the drug penetration coefficient, *θ_s_* we used a geometric sequence with ratio 0.55. Since we assumed full penetration of drug into blood (first compartment), *θ*_1_ = 1. For the compartments s = 2:10 we have: *θ_s_* = (0.55)^*s*-1^.

**Table 1:** Parameters.

| Definition | Units | Value | Ref |
| --- | --- | --- | --- |
| Target cell production rate ($\log_{10}(\lambda)$) | $\log_{10}(\text{Cells}/(\mu\text{l} \cdot \text{day}))$ | (1.54, 2.88) | [22] |
| Target cells death rate ($\log_{10}(dT)$) | $\log_{10}(1/\text{day})$ | (-1.35, -0.34) | [22] |
| Density dependent infection rate ($\log_{10}(\beta)$) | $\log_{10}(\text{ml}/\text{day})$ | (-5.78, -5.23) | [22] |
| Infected cell death rate ($\log_{10}(dI)$) | $\log_{10}(1/\text{day})$ | (-0.76, 0.42) | [22] |
| Virus production rate ($\log_{10}(\gamma)$) | $\log_{10}(1/\text{day})$ | (3.39, 4.00) | [22] |
| Virus clearance rate ($\log_{10}(dv)$) | $\log_{10}(1/\text{day})$ | 18.8 | [22] |
| Efficacy of drug ($\log_{10}(\eta)$) | --- | (0.60, 0.89) | [22] |
| Latently infected cell death rate (d) | 1/day | 0.004 | [7] |
| Latently infected cell proliferation rate ($\rho$) | 1/day | 0.2045 | [7] |
| Fraction of newly infected cells that turn into latently infected cell ($\alpha$) | --- | $10^{-6}$ | [7] |

## 3. Results

### 3.1. Dynamic of viruses, infected T-cells, and targeted T-cells

To investigate the change of viruses and infected cells, we ran a simulation where the treatment starts in the fifth year. To take into the account the effect of the tissue volume, we considered three volumes: 1ml, 30ml, and 250ml which represent, a single lymph node, a fraction of the body’s lymph nodes, and the total lymph nodes in the body, respectively (figure 4). As shown in figure 5 and figure 6, after ARV intensification, concentrations of viruses and infected cells in the blood drop more than the concentrations in the lymph node. When infection starts the concentration of T-cells as shown in figure 7 drops, but after administration of ARVs, the concentration of T-cells increases almost to the initial level. Infected cell and virus concentrations increase in the blood when the infected volume of the tissue increases. For the tissue, changing the volume from 30ml to 250ml does not change the concentrations, however, for the single lymph node the concentration is lower than in either the 30ml or 250ml cases.

**Figure 4:**
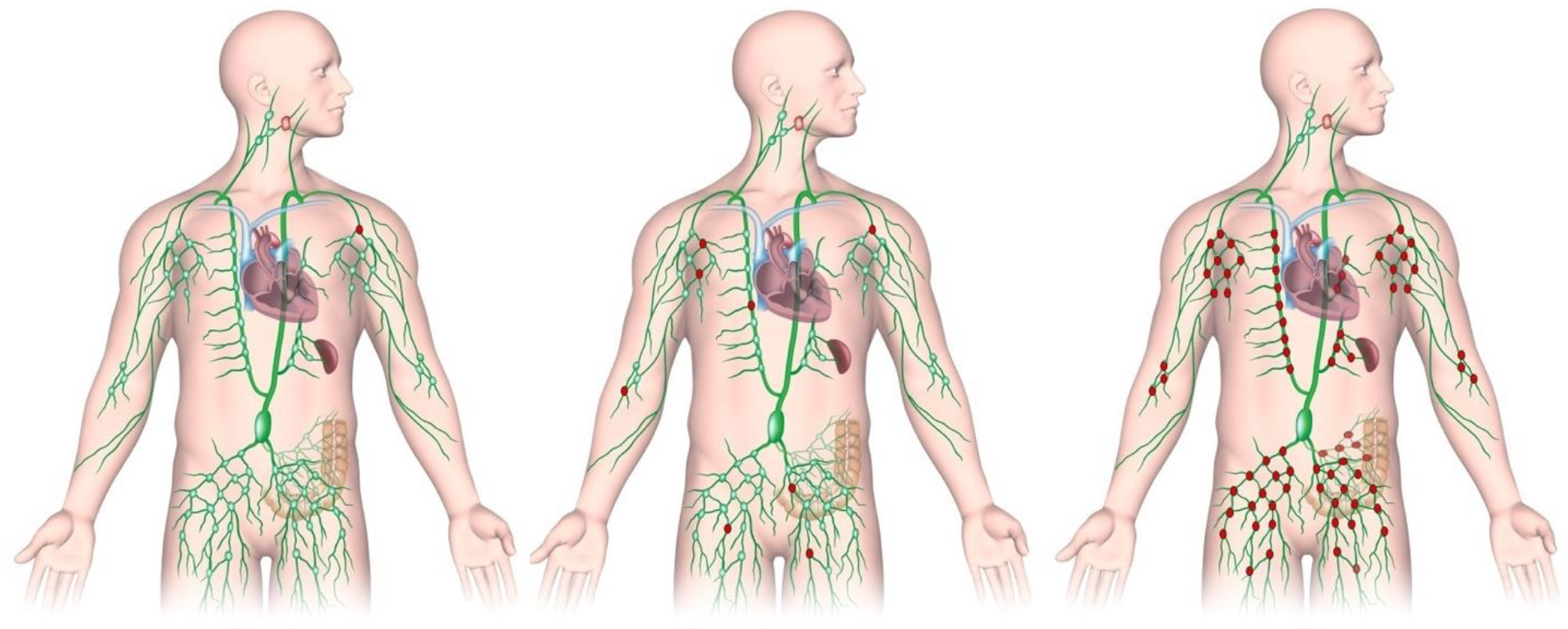
Illustration of lymphatic system. The red LNs shows LNs with active sanctuary site for ongoing replication. Right figure represents a single LN as active sanctuary site, the middle figure shows partial LNs as active sanctuary site, and the left figure shows the whole lymphatic system as active sanctuary site.

**Figure 5:**
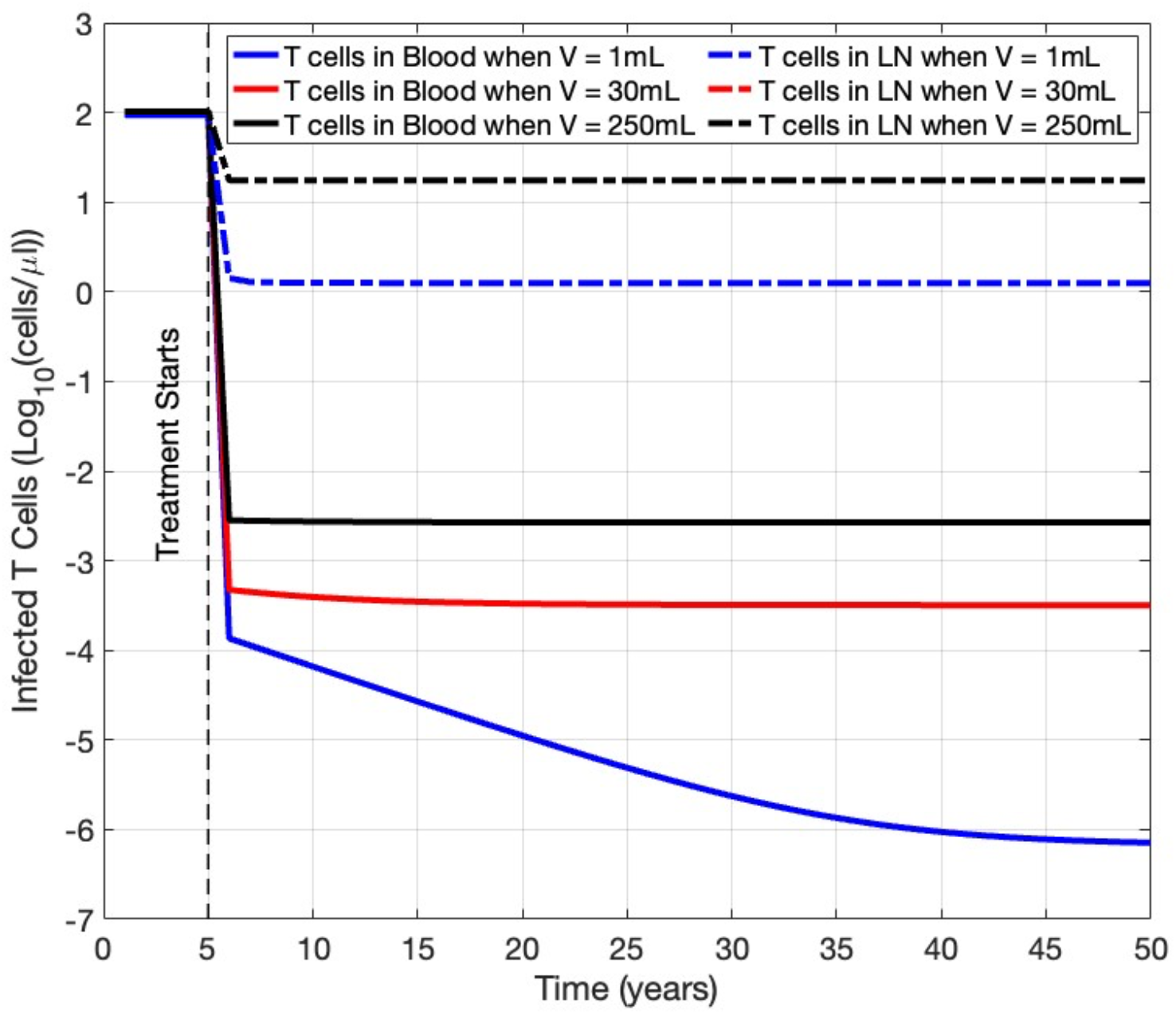
The dynamics of actively infected HIV cells in the blood and LN of patients with LN sanctuary sites of different sizes. Solid lines represents blood concentrations, dashed lines represent LN concentration. Blue lines show 1 ml sanctuary site, red lines show 30 ml of active sanctuary site and black lines show 250 ml or full lymphatic involvement of sanctuary site

**Figure 6:**
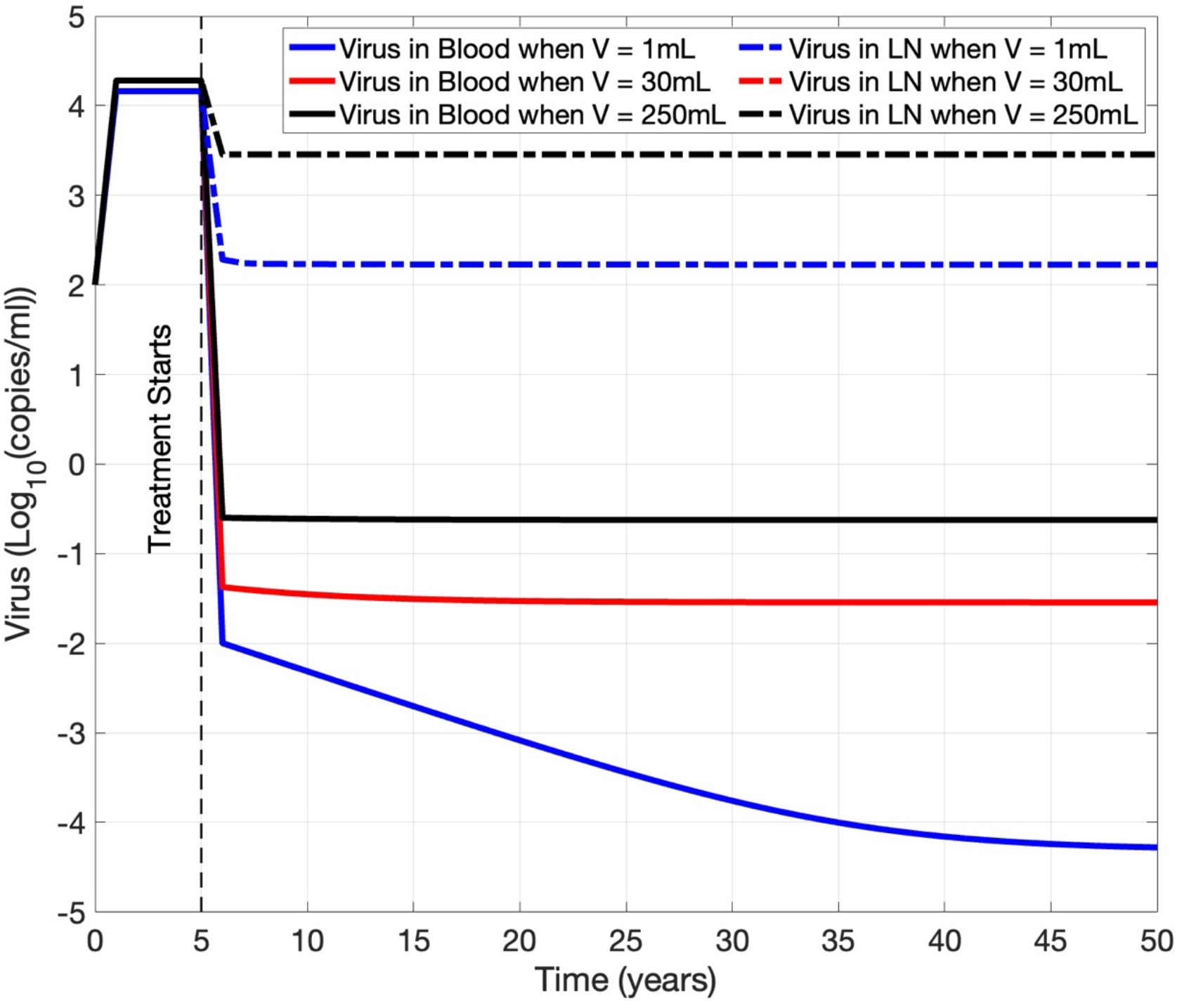
The dynamics of HIV viruses in the blood and LN of patients with LN sanctuary sites of different sizes. Solid lines represent blood concentrations, dashed lines represent LN concentration. Blue lines show 1 ml sanctuary site, red lines show 30 ml of active sanctuary site and black lines show 250 ml or full lymphatic involvement of sanctuary site.

**Figure 7:**
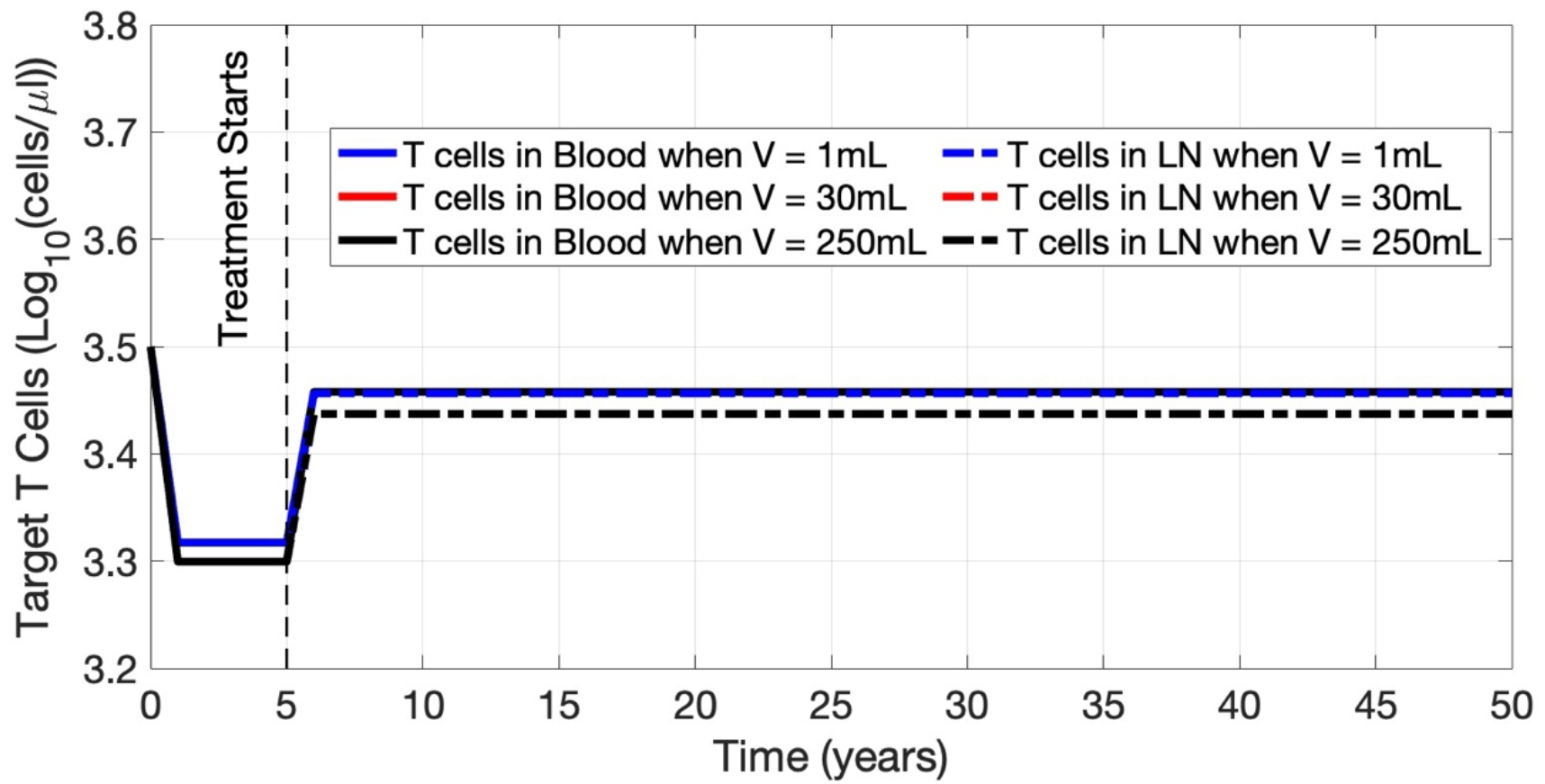
The dynamics of T-cells in the blood and LN of patients with LN sanctuary sites of different sizes. Solid lines represents blood concentrations, dashed lines represent LN concentration. Blue lines show 1 ml sanctuary site, red lines show 30 ml of active sanctuary site and black lines show 250 ml or full lymphatic involvement of sanctuary site.

### 3.2. Archival latent cells formation in sanctuary sites

To better understand the formation of novel latent cells after ARV intensification, we used a treatment plan that starts at the fifth year after infection and continues for the next 45 years. The parameter values were as described in table 1, and we assumed 30ml for the infected volume of the lymph nodes. The solid lines, and dashed lines in figure 8 show the concentration of latent cells in the blood and lymph node, respectively. As shown, the concentration of AL cells is equal in both compartments, while the concentration of NL cells is higher in the lymph nodes compared to blood. After the formation of NL cells start, it takes 27 and 45 years for NL cells to reach to the same concentration as AL cells in the lymph node and blood, respectively.

**Figure 8:**
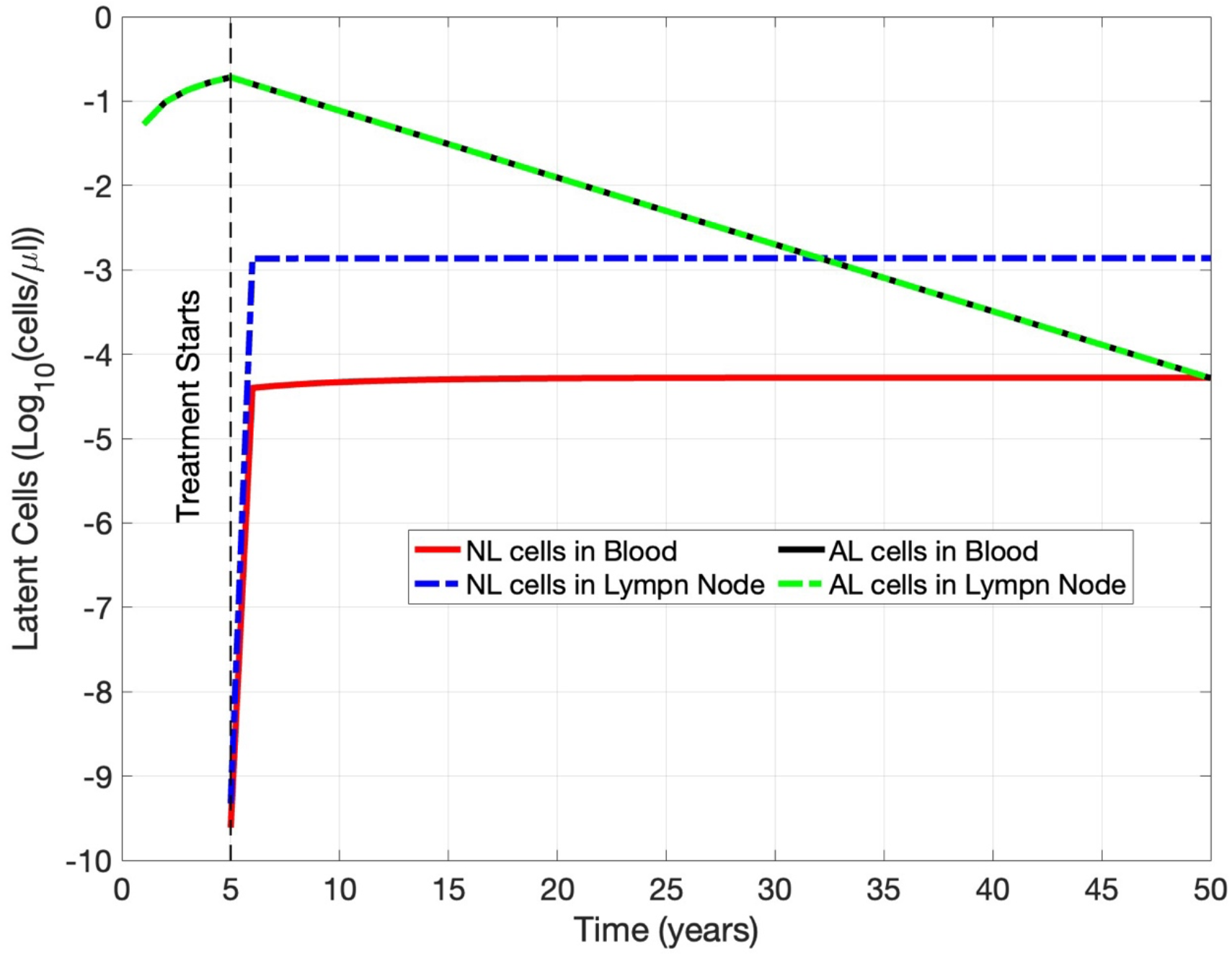
The dynamics of novel latent cells and archival latent cells in the blood and LN of patients with LN sanctuary sites. Solid lines represent blood concentrations, dashed lines represent LN concentration. Blue and red lines show NL cells, and black and green lines show AL cells.

### 3.3. Relation between sanctuary site volume and Archival latent cells formation

To understand the relationship between the volume of infected tissue and NL cell formation, we plotted the NL cell concentrations for both blood and lymph node compartments, while varying the infected lymph node volume between 1ml and 250ml which represent a single lymph node and all lymph nodes in the body, respectively. As shown in figure 9, when the infected volume increase, the concentration of NL cells increases in both blood and lymph node compartments, while the concentration in the tissue in higher than blood for all the volumes. It takes more time for NL cells to reach the steady state level inside the blood compared with lymph node.

**Figure 9:**
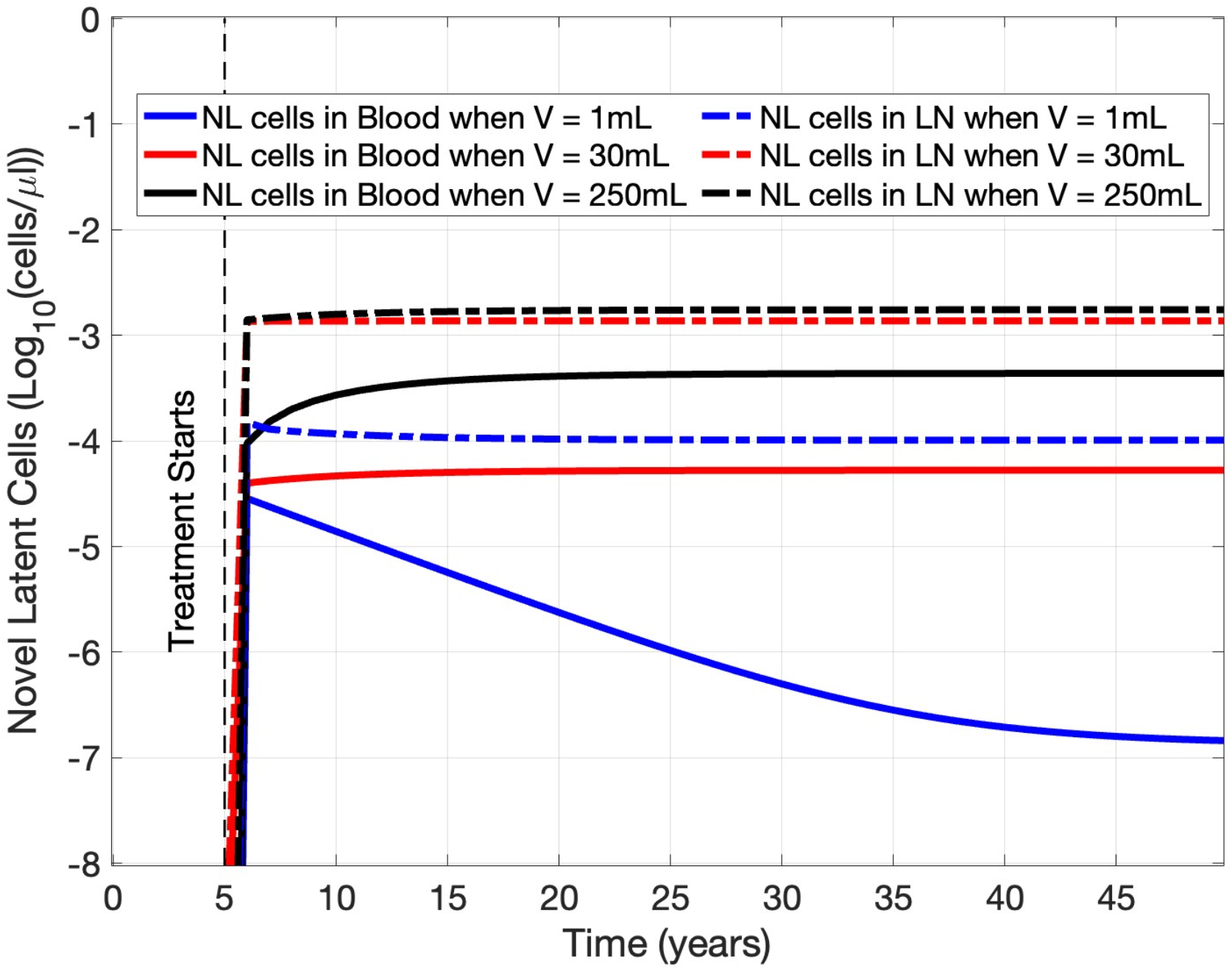
The dynamics of NL cells in the blood and LN of patients with LN sanctuary sites of different sizes. Solid lines represent blood concentrations, dashed lines represent LN concentration. Blue lines show 1 ml sanctuary site, red lines show 30 ml of active sanctuary site and black lines show 250 ml or full lymphatic involvement of sanctuary site.

### 3.4. Ratio of Novel latent cells over total latent cells

To better understand the time needed for novel latent cells to become dominant in the tissue and blood, we plotted the ratio of NL cells over the total latent cells. We started the treatment from the fifth year, and we considered three different volumes of sanctuary sites as shown in figure 10. For a realistic condition where 30 ml of lymph nodes were infected, it takes 33 years for NL cells to reach 10% of the total latent cells in the blood after treatment starts. This result illustrates why it is hard to capture NL cells in blood samples. Even inside the lymph nodes it takes 15 years for NL cells to reach the 10% level. For an extreme condition where all lymph nodes were infected it takes 22 and 13 years for NL cells to reach the 10% level inside the blood and tissue, respectively, as shown in figure 10 For a single lymph node even after 45 years the NL cells don’t reach 10% of the latent cell population in the blood.

**Figure 10:**
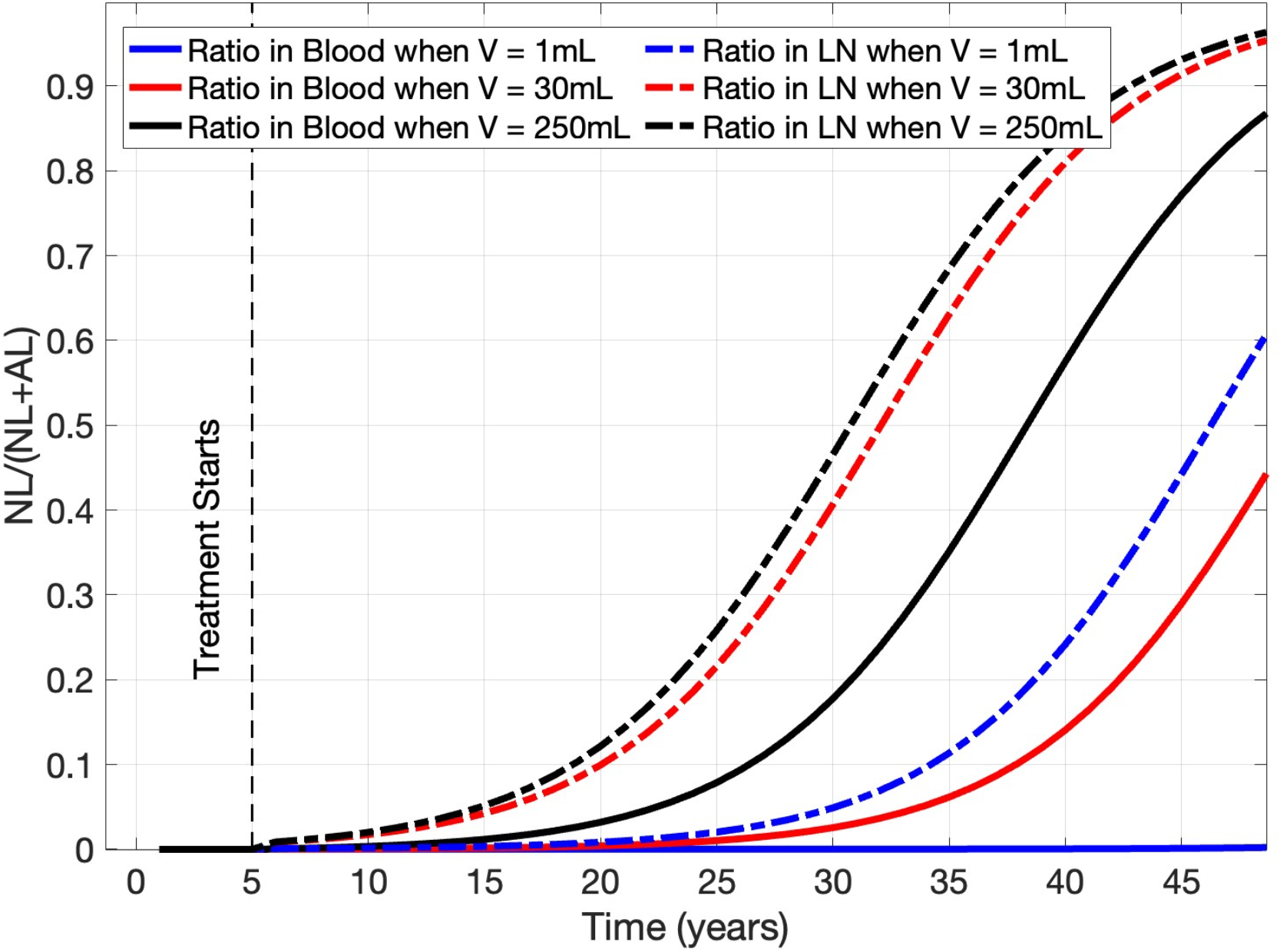
Ratio of NL cells over total latent cells in the blood and LN of patients with LN sanctuary sites of different sizes. Solid lines represent blood concentrations, dashed lines represent LN concentration. Blue lines show 1 ml sanctuary site, red lines show 30 ml of active sanctuary site and black lines show 250 ml or full lymphatic involvement of sanctuary site.

### 3.5. Probability of viral evolution occurring

To investigate the ratio of the NL cells in the samples analyzed by papers that did not detect viral evolution in HIV-1 sequences, we summarized the NL/(AL+NL) for all the samples in those papers [17, 23, 3] in table 2. We considered 30ml for partial volume of infected lymph nodes total lymph nodes infection which is an extreme condition. In [13, 14] they only examined samples from blood and the ratio of the NL cells to total cells varies among 0.15% to 1.79%, which are very small ratios. In [23, 3], authors analyzed samples from both blood and lymph nodes. In [23] the NL/(AL+NL) is between 0.37-0.94% for blood and 3.67-4.44% in lymph nodes. For [3] because of the long observation time, the ratios were higher than other papers, and it was 6.74% for the blood and 12.39% for the lymph nodes. But, we need to keep this fact in mind that not all those lymph nodes in the body is a center for ongoing replications so despite the high ratio of the NL cells it does not guarantee to detect ongoing replication. In table 2 based on the number of HIV-1 sequences analyzed for each patient, and the ratio of the NL cells to total latent cells, the probability of detecting at least one NL cell is calculated (p). In [17, 18] the probability is varying below the 37% which demonstrate the low chance of getting even one NL cells in the blood in these studies. In [23, 3], the probability of detecting at least one NL cells in the lymph node are more than 47%. As we discussed before the chance of choosing the right lymph node is also low and we know one sample is not enough to detect the viral divergent.

**Table 2:** Model prediction of Probability of detecting at least one NL cell.

| Papers | DI<sup>1</sup> before ART | Sampling Time<sup>2</sup> | NL/(NL+AL) | # of Seq. | P<sup>3</sup> |
| --- | --- | --- | --- | --- | --- |
| [17] | 5 years | 12 years | B<sup>4</sup>: 0.41% | 27 | 1.21% |
| [17] | 5 years | 4.2 years | B: 0.09% | 22 | 0.21% |
| [17] | 5 years | 10 years | B: 0.27% | 24 | 0.75% |
| [17] | 5 years | 9.4 years | B: 0.25% | 4 | 0.09% |
| [17] | 5 years | 8.8 years | B: 0.22% | 3 | 0.05% |
| [23] | 1 year | 1.8 years | B: 0.24%<br>T<sup>5</sup>: 7.67% | B: 54<br>T: 18 | B: 1.43%<br>T: 8.9% |
| [23] | 2 years | 5.7 years | B: 0.41%<br>T: 0.34% | B: 37<br>T: 26 | B: 0.18%<br>T: 0.98% |
| [23] | 4.3 years | 8.5 years | B: 0.41%<br>T: 0.31% | B: 32<br>T: 40 | B: 0.11%<br>T: 1.36% |
| [3] | 5 years | 19.1 years | B: 0.88% | 46 | 3.95% |
| [3] | 5 years | 22.7 years | B: 31.06% | 20 | 11.99% |
| [3] | 5 years | 22.7 years | B: 31.06% | 30 | 12.00% |

1: Duration of infection
2: Sampling time after ART
4: Sampling from the blood
5: Sampling from the tissue

### 3.6. Detecting viruses and latent cells in different generations

A phylogenetic tree is a branching diagram showing the evolutionary relationships among various species. The pattern of branching in a phylogenetic tree represents how species evolved from a common ancestor. Phylogenetic trees are used to detect viral ongoing replication in HIV patients [2, 34]. Figure 11 shows phylogenetic tree for two different sample point of a patient. *b*_1:10_ represent samples obtained at first time point and *h*_1:10_ represent samples obtained at second time point. To go from sequence *a*_6_ to *b*_0_ it requires 7 point mutations, implying this could not happen without seven intervening sequential infection to accumulate seven necessary mutations. That is why counting not just the total number of NL cells but how many generations exist, is important. In this study a model is developed (Appendix A) to measure the viruses and associated infected cells in different generations, in order to track the number of viruses in each generation. To keep the model tractable, we make the last generation of viruses the accumulative generation of 10th generation and higher. As shown in figure 12 and figure 13, the number of NL cells and infected cells in higher generations are proportional to total number of total NL cells at each time point, so the results derived from tracking all NL cells represent an upper bound on the detectability of ongoing replication through phylogenetic analysis.

**Figure 11:**
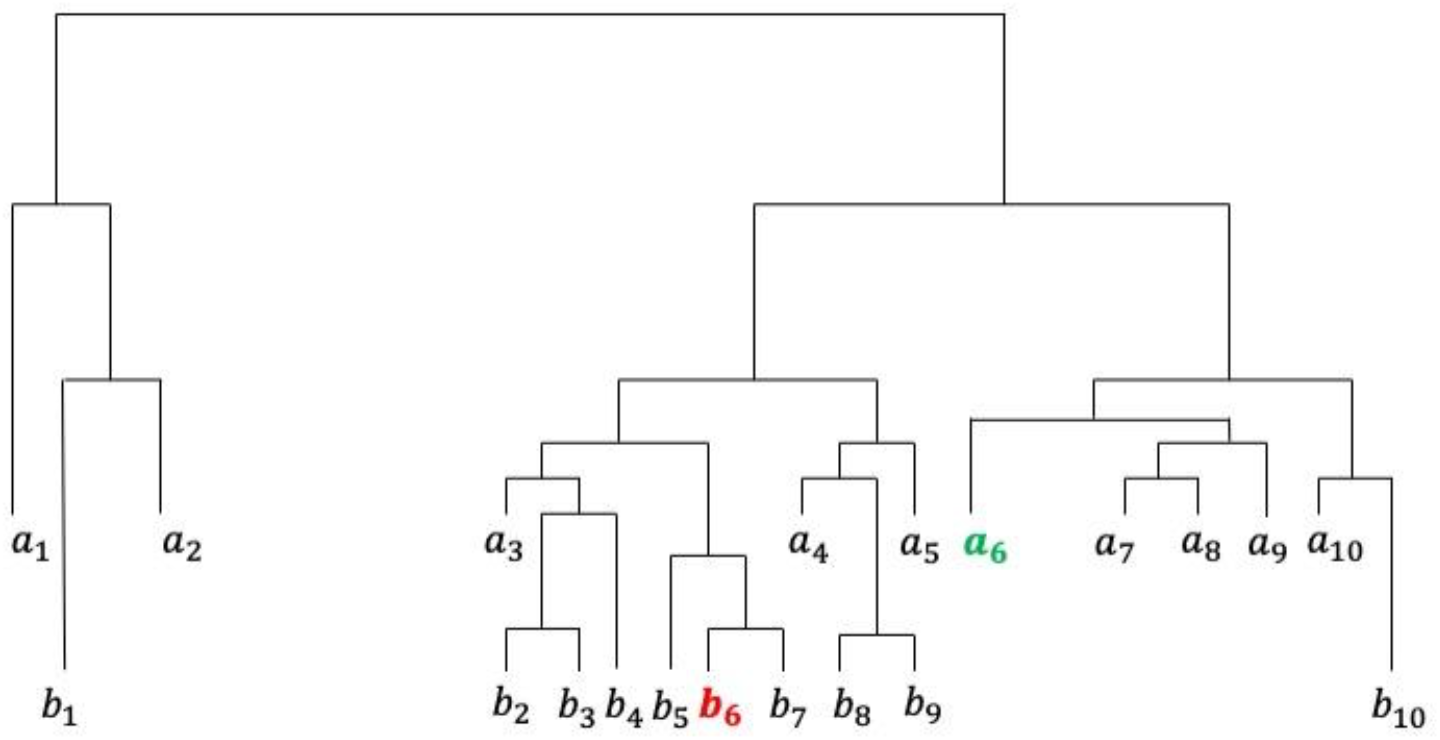
A phylogenetic tree where a1:10 and b1:10 show the samples obtained at different time points.

**Figure 12:**
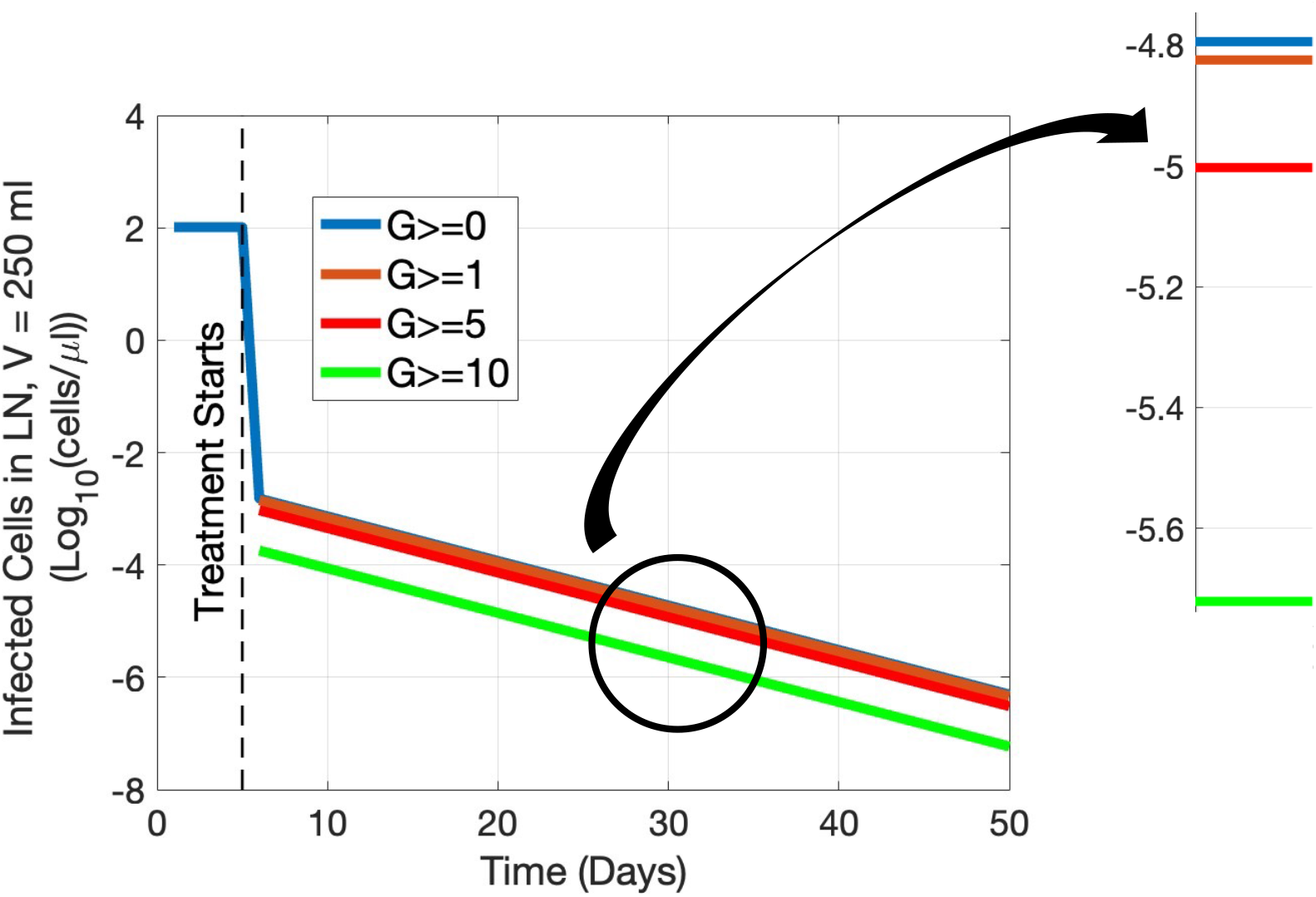
The dynamics of NL cells in the LN for different generations of the infected cells. Blue line shows generation of zero and more, orange line shows generation of 1 and more, red line shows generation of 5 and more, the green line shows generation of 10 and more.

**Figure 13:**
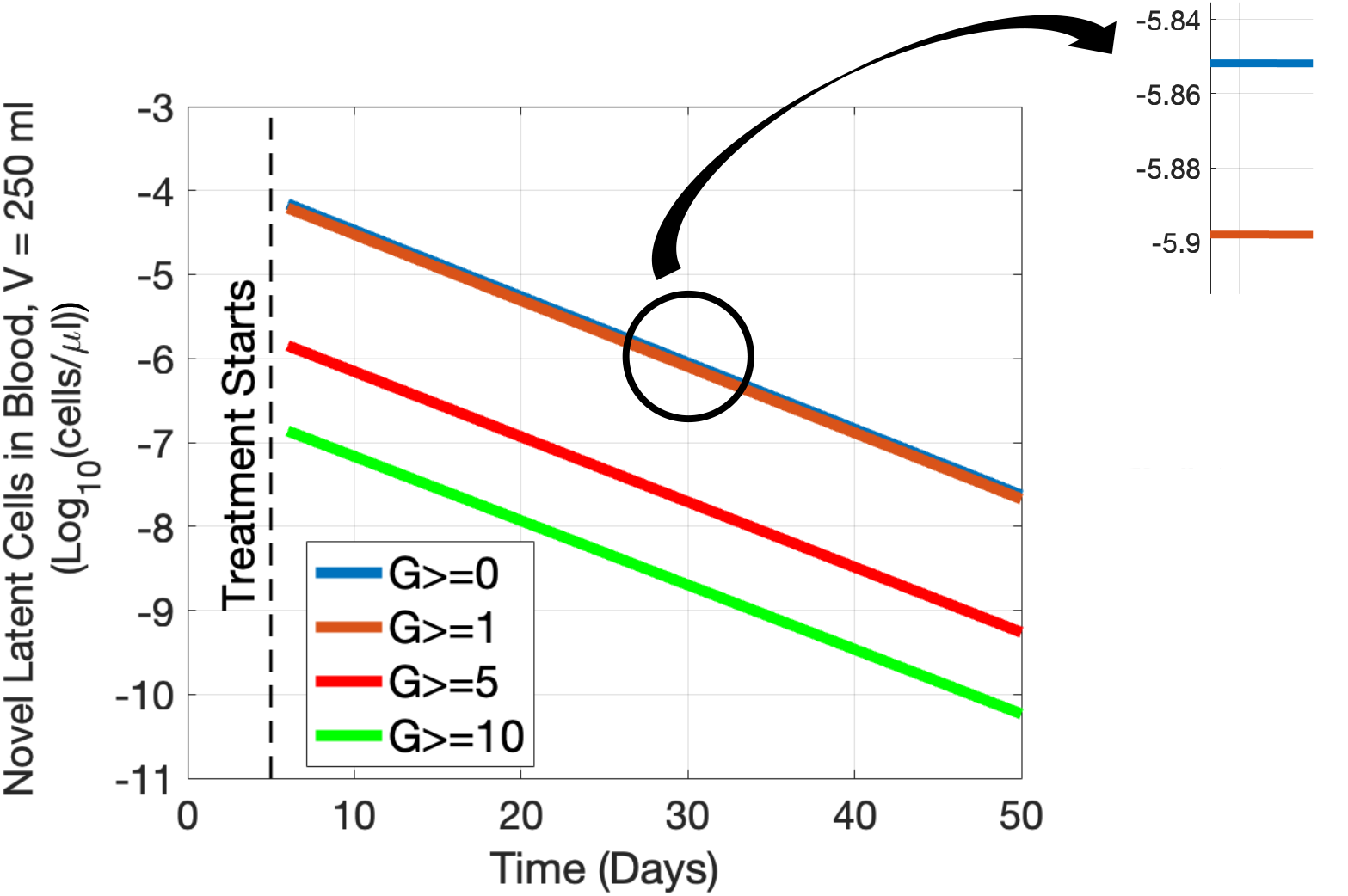
The dynamics of actively infected HIV cells in the LN for different generations of the infected cells. Blue line shows generation of zero and more, orange line shows generation of 1 and more, red line shows generation of 5 and more, the green line shows generation of 10 and more.

## 4. Discussion

The existence of ongoing HIV viral replication in sanctuary sites is a controversial topic. Phylogenetic analysis of virus samples from the blood has mostly failed to show any evidence of ongoing replication. This study shows that detecting ongoing replication via phylogenetic divergence from blood samples depends on the ability to sample a significant number of latent cells infected after the initiation of therapy. We developed a mathematical model that tracks replication after treatment in patients with ongoing replication in lymph node sanctuary sites.

Our results show that if a single lymph node with a volume of 1ml were to act as an isolated sanctuary site, according to our model, after 50 years, there would still be no measurable quantity of NL cells circulating in the blood. On the other hand, if the entire lymphatic system, which has a total volume of 250ml, behaves as a sanctuary site, it would take 22 years before NL cells reach 10% of the entire circulating latent cell population. If you sample at any time point, you need to acquire a certain number of NL cells; those NL cells also have to contain mutations genetically divergent from the existing phylogeny, which is the worst-case scenario. Despite considerable levels of ongoing replication in the lymphatics, we would nevertheless need to wait 22 years to see 10% of the circulating latent cells being NL cells, and even if we sequence a subset of those NL cells, we may not get enough information to conclude that we have ongoing replication.

The odds of detecting ongoing replication improve significantly if we biopsy lymph nodes directly instead of sampling the blood, as the NL cells appear to remain largely compartmentalized after formation. But this runs into a separate issue; not all the lymph nodes are necessarily acting as sanctuary sites, and a biopsy of an uninvolved lymph node will not contain any more NL cells than a biopsy of the blood. Nevertheless, we have simulated the percentage of latent cells within involved LNs over time and have shown that the chance of detecting at least one NL cell is not more than 12%.

Several other papers have shown no evidence of viral divergence [17, 23, 3]. We have simulated the conditions from these various studies, and by calculating the probability of detecting at least one NL cell, we showed that these studies would have been unlikely to detect viral divergence even if these patients had active lymph node sanctuary sites. In [17], Kearney et al., by analyzing chronically infected patients who were under cART treatment for up to 12 years, showed that, regardless of significant changes in population structure, the genetic sequence of the virus after long-term cART is not significantly different from viruses before cART. Five patients were sampled on long cART after 4.2 to 12 years. 3 to 27 samples were gathered from each patient. Our model calculated the probability of detecting at least one NL cell based on the sampling time and number of sequences. The probability varies between 0.05% to 1.21%, which are very small numbers to guarantee detecting at least one NL cell, and the probability of detecting sufficiently divergent mutations is even smaller.

McManus et al. [23], investigated the possibility of viral replication in lymph nodes and examined the HIV-1 sequence from 5 patients after up to 13 years on cART. They found identical viral sequences both in lymph nodes and blood during cART, and no evidence of significant divergence from the pre-cART viral sequence was detected either in a lymph node or blood. Samples from the HIV-1 infected patients were obtained after 1.8, 5.7, and 8.5 years of treatment. 18 to 54 sequences were gathered among the patients. Our model calculated the probability of detecting at least one NL cell from 0.11% to 1.43% in blood and 0.98% to 8.9% in the lymph node.

Other research by Bozzi et al. [3] has been conducted to investigate the ongoing replication of HIV-1 in sanctuary sites such as lymph nodes with low ART penetration. They analyzed samples from patients under cART for 8 to 16 years. They did not detect any sequential divergence among samples, and they found that the clonal expansion of infected cells before cART treatment is responsible for maintaining HIV during treatment. They sampled 20 to 46 sequences from HIV-1 patients under cART after 19.1 and 22.7 years from blood and lymph nodes. Our model detected a 12% and 3.95% chance of detecting at least one NL cell in the LN and blood, respectively.

Our results showed that for patients with as many as all of their lymph nodes involved in ongoing replication as sanctuary sites, it takes 22 years to accumulate a detectable quantity of novel latent cells. Furthermore, merely detecting a novel latent cell would not be sufficient to determine that it was a novel latent cell. An adequate amount of genetic divergence from the preexisting sequences must be existed to determine the novel latent cell. When we augment the model to track the generations of the virus in those novel latent cells, the odds get even worse for detecting novel latent cells from higher generations.

This harmonizes the previous findings that show evidence of ongoing replication in sanctuary sites, with the previous studies showing no evidence of genetic divergence in patients on long-term treatment. It shows that those methods for detecting divergence are not sensitive enough to see the small amount of replication in isolated sites.

## 5. Appendix

In this section the model that used to describe the dynamics of viruses and the immune system based on their generation. *v_s,g_* represents the gth generation of the virus in compartment s. *u*_*s,g*=1_ is the initial virus that infect the body. Equation 11 describes the infection of T-cells by viruses of all generations.

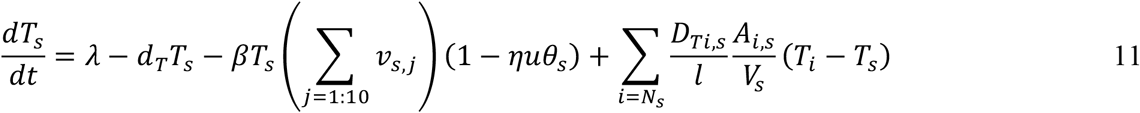

Depending on which generation of viruses infected by that determines the generation of the latent infected T-cells and infected T-cells shown I equations 12–14.

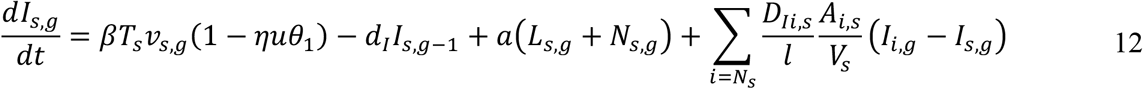

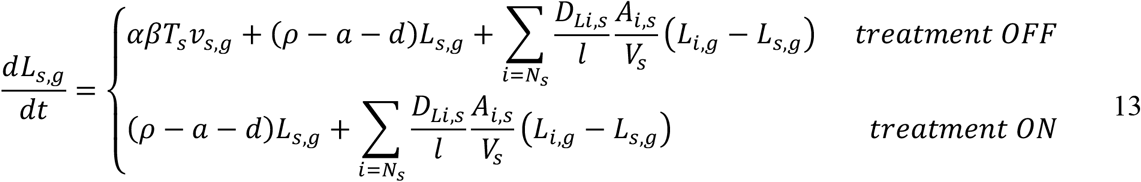

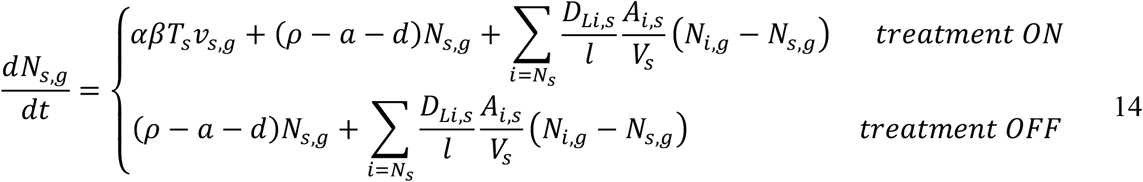

Viruses that released into the body by the ith generation of infected cells create the (i+1)th generation of viruses as shown in equation 15.

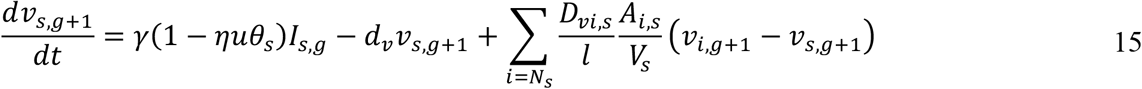

To keep the model tractable, we make the last generation of viruses the accumulative generation of 10th generation and higher *v*_*s,g*=10_ = *v*_*s,g*=10_ + *v*_*s,g*=1_.

